# A foldable respiratory siphon enables near-surface locomotion through asymmetric rowing in *Helophilus* larvae

**DOI:** 10.64898/2026.06.29.735254

**Authors:** Teruyuki Matsunaga, Akinao Nose

**Affiliations:** Department of Physics, Graduate School of Science, The University of Tokyo, Tokyo, Japan; Department of Complexity Science and Engineering, Graduate School of Frontier Sciences, The University of Tokyo, Chiba, Japan

**Keywords:** Hoverfly larvae, respiratory siphon, folding–unfolding, aquatic locomotion, biomechanics, multifunctionality

## Abstract

Specialized animal structures often perform multiple functions, yet how their structural adaptations support distinct behaviors remains poorly understood. Eristaline hoverfly larvae, commonly known as rat-tailed maggots, possess exceptionally elongated and extensible posterior respiratory siphons that allow them to breathe while submerged. Here, we investigated the structural basis of siphon extension and its potential locomotor function in *Helophilus virgatus* larvae. Fluorescence labeling showed that siphon elongation involves the unfolding of previously concealed structures, while phalloidin staining revealed an extensive array of transversely oriented muscle fibers within the siphon. Behavioral observations further revealed a previously undescribed mode of horizontal locomotion just beneath the air–water interface, involving asymmetric rowing movements of the posterior siphon. Kinematic analyses showed that most forward body displacements occurred during the backward stroke, which was associated with increased siphon straightness, whereas the forward stroke produced little net forward displacement. These findings reveal structural features underlying siphon extensibility and demonstrate that a specialized respiratory organ also contributes to locomotion, highlighting the multifunctionality of specialized body structures in aquatic environments.

## Main

Animals inhabiting aquatic environments face physical challenges associated with oxygen availability and locomotion through water [1]. Adaptations to these challenges often involve specialized body structures that interact mechanically with the surrounding environment [2]. For example, some aquatic beetles possess plastrons that facilitate underwater respiration [3]. Moreover, dragonfly nymphs use their rectal chamber for both respiration and jet-propelled locomotion[4,5]. These examples illustrate diverse adaptations to aquatic environments, including the use of a single organ system for multiple functions. However, the extent to which specialized respiratory structures are also employed for locomotion remains poorly understood.

Hoverfly larvae (Diptera: Syrphidae) provide an opportunity to examine how specialized respiratory structures can serve additional functions. Although adult hoverflies are important pollinators, their larvae occupy remarkably diverse ecological niches, ranging from terrestrial predators to aquatic saprophages [6–10]. Among Eristalinae, larvae of several lineages inhabit stagnant ponds, marshes, and other aquatic environments rich in organic matter and often depleted in dissolved oxygen [11,12]. These aquatic larvae, commonly known as rat-tailed maggots, possess an exceptionally elongated and extensible posterior respiratory siphon [11,13,14] (Fig. 1a). By extending the siphon to maintain contact with the air–water interface while remaining submerged, larvae can obtain atmospheric oxygen without repeatedly surfacing [15].

**Figure 1.**
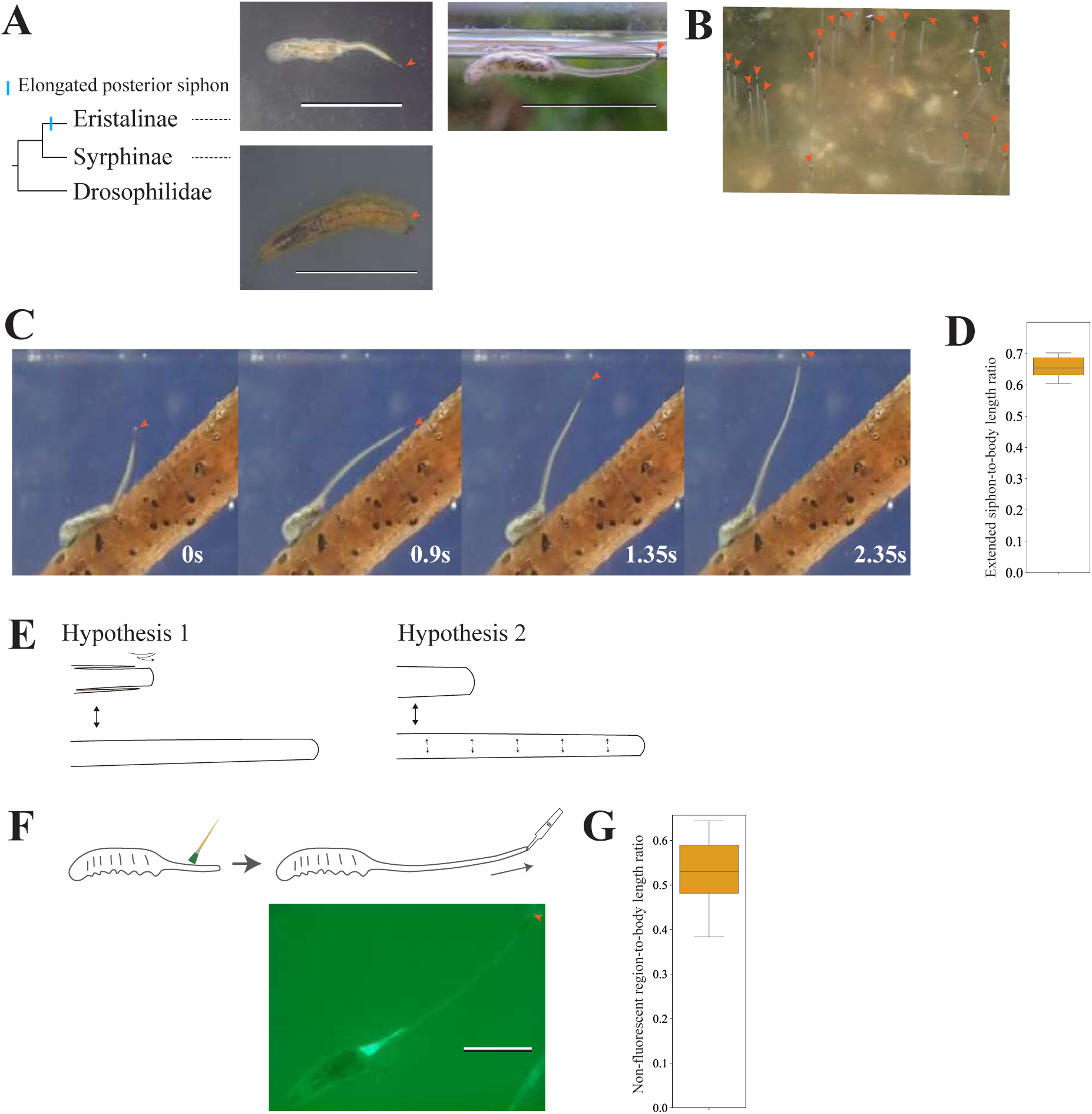
*Helophilus* larvae (rat-tailed maggots) use an extensible posterior siphon for respiration. **(a)** Schematic phylogeny showing Syrphidae (including Eristalini and Syrphini) and Drosophilidae as an outgroup (branch lengths not to scale). Representative dorsal views of *Helophilus virgatus* and *Episyrphus* sp. larvae on agar are shown on the left (scale bars, 0.5 cm and 1 cm, respectively). The *Episyrphus* sp. larva is shown with aphid prey remains attached near the posterior end. The right panel shows an *H. virgatus* larva floating on the water surface. Orange arrowheads indicate the posterior spiracles. **(b)** Natural habitat of *H. virgatus* larvae aggregating in shallow water with their posterior siphons extended to maintain contact with the air–water interface for respiration. **(c)** Time-lapse images showing sequential extension of the posterior siphon as an *H. virgatus* larva descends beneath the water surface while maintaining contact with the air–water interface. **(d)** Box plot showing the extended posterior siphon-to-body length ratio of *H. virgatus* larvae (*n =* 10). **(e)** Schematic illustration of two proposed mechanisms of posterior siphon extension: folding–unfolding (left) and tissue expansion (right). **(f)** Fluorescent labeling assay to distinguish between the two mechanisms. The upper panel illustrates the experimental procedure, in which the posterior siphon was coated with fluorescent dye in its contracted state and subsequently extended manually. The lower panel shows a representative posterior siphon after manual extension. A previously unlabeled region became exposed in the distal portion of the siphon, consistent with the unfolding of a previously concealed region. The posterior end is indicated by an orange arrowhead. Scale bar 1 cm. **(g)** Box plot showing the ratio of the non-fluorescent region length to body length (*n* = 6 larvae).

The remarkable extensibility of the posterior respiratory siphon has been attributed to the unfolding of folded anal segments [16]. Moreover, beyond respiration, eristaline larvae can regulate their buoyancy by altering the volume of air within their respiratory system [17]. However, the structural changes accompanying siphon extension and the organization of its associated musculature remain poorly characterized. Furthermore, whether movements of the posterior siphon itself contribute to horizontal locomotion remains unknown. Examining both siphon structure and movement kinematics may therefore reveal how a specialized respiratory organ supports an additional locomotor function.

Here, we investigated the structure and function of the extensible posterior respiratory siphon in larvae of the hoverfly *Helophilus virgatus* (Eristalinae). Using behavioral observations, fluorescence labeling, morphometric analyses and phalloidin staining, we provide experimental evidence that siphon extension involves the unfolding of folded anal structures and characterize its distinctive transverse muscle architecture. We further describe a previously undocumented mode of horizontal locomotion just beneath the air–water interface, involving asymmetric rowing movements of the posterior siphon. Kinematic analyses reveal that most forward body displacement occurs during the backward stroke, when the siphon is relatively straight, whereas the forward stroke produces little net forward displacement. Together, these findings demonstrate that a specialized respiratory organ can also contribute to locomotion, highlighting the multifunctionality of specialized body structures in aquatic environments.

## MATERIAL AND METHODS

### Sampe collection and rearing

Larvae of *Helophilus virgatus* were collected from a small stagnant pond in Nagareyama, Chiba, Japan (35.905°N, 139.905°E), in November 2024, April 2025, and May 2026. Larvae of *Episyrphus* sp. were collected from *Brassica* leaves in Yaita, Tochigi, Japan (36.851°N, 139.889°E), in November 2025. *H. virgatus* was identified by cytochrome c oxidase subunit I (COI) sequencing and confirmed by morphological characteristics. *Episyrphus* sp. was identified based on morphological characteristics. COI was amplified using primers described previously [18] and sequenced using Sanger sequencing. A Canton-S strain of *Drosophila melanogaster* was obtained from the Kyoto Stock Center. All species were maintained at 25°C under a relative humidity of >60%. *D. melanogaster* was reared on standard cornmeal medium. *H. virgatus* larvae were maintained in water collected from the original habitat. *Episyrphus* sp. larvae were reared on fresh *Brassica* leaves collected from the original collection site.

### Visualization of folded posterior siphon structures

The protocol was adapted from a previous study [19]. Briefly, *H. virgatus* larvae were rinsed with distilled water and gently dried using Kimwipes. While the posterior siphon was in a contracted state, its surface was coated with a commercially available green fluorescent dye using a paintbrush. After 5–10 s, the posterior siphon was manually extended using forceps. Images were acquired using a stereomicroscope (Olympus SZX16) equipped with a digital camera (Hozan L-836). Representative still images extracted from the videos were used for analysis. The non-fluorescent region length was defined as the length of the unlabeled portion of the posterior siphon exposed after extension. The non-fluorescent-to-body length ratio was calculated by dividing the non-fluorescent region length by the total body length, measured from the anterior tip to the posterior tip.

### Body wall muscles staining and counting of muscle fibers

The protocol was adapted from a previous study [20]. Briefly, larvae were dissected in phosphate-buffered saline (PBS) to prepare fillet preparations on Sylgard-coated dishes (Sylgard 184, Dow) using fine forceps, microscissors (LMB-54-1, Nazme), and insect pins. Samples were fixed in 4% formaldehyde for 15 min and subsequently rinsed in PBT (PBS containing 0.2% Triton X-100). Samples were then blocked in 10% normal goat serum in PBT and incubated overnight at 4°C with FITC-conjugated phalloidin (1:100 dilution in blocking solution). After rinsing in PBT, larval musculature was imaged using a confocal microscope (FV3000, Olympus) equipped with a 488-nm laser and either a 4× air objective (UPLSAPO, Olympus) or a 20× objective (XLUMPlanFL N, Olympus).

For *H. virgatus*, low-magnification confocal images acquired using the 4× objective were stitched using the Pairwise Stitching plugin in ImageJ [21]. Muscle fiber orientations were quantified in ImageJ by measuring their angles relative to the longitudinal axis of the posterior siphon. In *H. virgatus*, measurements were performed on the left side of the posterior siphon, with the proximal, middle, and distal regions analyzed separately. In *Episyrphus* sp. and *D. melanogaster*, measurements were performed across the entire musculature of abdominal segment A8.

To quantify the abundance of transverse muscle fibers, we counted the number of transverse fibers intersecting the longitudinal axis of the posterior siphon in *H. virgatus* and a corresponding longitudinal reference line in abdominal segment A8 of *Episyrphus* sp. and *D. melanogaster*. Transverse muscle fibers were defined as fibers oriented at an angle of 90° ± 30° relative to the longitudinal axis.

### Measurement of dorsal trunk width

*H. virgatus* and *D. melanogaster* larvae were placed in a Petri dish filled with distilled water, with their posterior spiracles maintained at the air–water interface. Videos were acquired using a stereomicroscope (Olympus SZX16) equipped with a digital camera (Hozan L-836) at 30 fps for 1 min. Representative still frames extracted from the videos were analyzed using ImageJ. The dorsal trunk-to-body width ratio was calculated by dividing a dorsal trunk width by the body width. Both measurements were taken at the midpoint between the mouth hooks and the anus along the anterior–posterior body axis.

### Siphon length measurements and near-surface locomotion assays

For measurements of posterior siphon elongation, a branch of *Zelkova serrata* collected on campus was placed in a 5-cm cubic glass chamber containing water collected from the original habitat, with approximately half of the branch submerged. Posterior siphon extension was recorded for 5 min from the lateral view using a digital camera (Olympus TG-6) in macro mode. The posterior siphon-to-body length ratio was calculated by dividing the maximum posterior siphon length by the body length. Body length was measured from the anterior tip to the base of the posterior siphon.

For locomotor analyses in Fig. 3, larval movements just beneath the air–water interface were recorded from a lateral view using the same imaging setup without a branch. Body size (cm) was defined as the body length measured from the anterior tip to the posterior end of the body when the posterior siphon was fully contracted. The rowing cycle period (s) was defined as the duration of one complete rowing cycle of the posterior siphon. Body displacement per cycle (cm) was defined as the net horizontal displacement of the midbody point between the beginning and end of a rowing cycle.

Swimming speed (cm s⁻¹) was calculated by dividing body displacement per cycle by the rowing cycle period. Normalized body displacement was calculated by dividing body displacement per cycle by body size.

For detailed kinematic analyses in Fig. 4, videos were recorded at 60 frames s⁻¹, and the positions of the posterior siphon tip, siphon base and anterior tip of the larval body were tracked throughout a single rowing cycle. The midbody point was defined as the midpoint between the anterior tip and the siphon base and was calculated for each frame. The trajectory of the siphon tip relative to its base was calculated by subtracting the coordinates of the siphon base from those of the siphon tip at each frame. The initial relative position was then subtracted so that each trajectory began at the origin (0, 0). Displacement of the midbody point along the x-axis was calculated relative to its position at the beginning of the cycle. The positive x- and y-directions were defined as leftward (forward) and downward, respectively.

The base–tip angle (θ) was defined as the angle between the horizontal water surface and the line connecting the siphon base to its tip. Siphon straightness was calculated for each frame as the straight-line distance between the siphon base and tip divided by the contour length of the siphon measured in the same frame. The unwrapped base–tip angle (θ) was smoothed using a centered three-point moving average, and angular velocity was then calculated using numerical differentiation with respect to time. For visualization, siphon tip trajectories were also smoothed using a three-point moving average.

A rowing cycle was defined as the interval between two successive instances when the posterior siphon tip reached its highest vertical position. Each rowing cycle was divided into two phases based on the transition of angular velocity from positive to negative. The transition time was estimated by linear interpolation between consecutive angular velocity measurements surrounding the zero crossing. Phase I (forward stroke) extended from the beginning of the rowing cycle to this transition, whereas Phase II (backward stroke) extended from the transition to the end of the cycle. Midbody displacement along the x-axis was normalized by the net displacement over the entire rowing cycle, such that the values at the beginning and end of the cycle were 0 and 1, respectively. The midbody position at the phase transition was estimated by linear interpolation. Displacement during each phase was calculated as the difference in midbody position between the beginning and end of that phase and was normalized by the net midbody displacement over the entire cycle.

Kinematic rowing effectiveness was defined as the absolute net displacement of the midbody point along the x-axis divided by the total path length traveled by the posterior siphon tip relative to its base during a single rowing cycle.

### Statistical analysis

For the analyses presented in Fig. 3, differences between groups were assessed using Mann– Whitney U tests, and correlations were evaluated using Pearson’s correlation coefficients (*r*). Differences between the two phases of the rowing cycle in Fig. 4d were assessed using a two-sided Wilcoxon signed-rank test. Box plots show medians and interquartile ranges (IQRs), with whiskers extending to the most extreme data points within 1.5 × IQR of the lower and upper quartiles, unless otherwise indicated. In Fig. 4, time-series data are presented as means ± standard errors of the mean (SEMs) for visualization. All statistical analyses were performed using Python.

## RESULTS

### *Helophilus* larvae possess an exceptionally elongated posterior respiratory siphon

Aquatic larvae of *Helophilus virgatus* were collected from a stagnant pond in Chiba, Japan. In their natural habitat, larvae were observed on floating materials such as branches, at the water surface, and aggregating in shallow water (Fig. 1a, b). Consistent with previous descriptions of Eristalinae larvae, *H. virgatus* possessed an exceptionally extended posterior respiratory siphon [13]. In contrast, terrestrial *Episyrphus* sp. larvae collected from *Brassica* leaves in Tochigi, Japan, and *Drosophila melanogaster* larvae lacked an elongated posterior siphon (Fig. 1a).

To quantify posterior siphon extension, *H. virgatus* larvae were placed in a water-filled chamber containing a partially submerged branch. Larvae frequently crawled beneath the water surface while maintaining contact with the air–water interface by extending their posterior siphons (Movie S1; Fig. 1c). The posterior siphon reached a median length corresponding to 65.4% of the body length (median posterior siphon-to-body length ratio = 0.654, IQR = 0.628–0.686, *n* = 10; Fig. 1d).

### Unfolding of concealed structures contributes to posterior siphon extension

The remarkable extensibility of the posterior respiratory siphon has previously been attributed to the unfolding of folded anal segments [16]. To experimentally examine this proposed mechanism in *H. virgatus* larvae, we considered two possible mechanisms of siphon extension: the unfolding of previously concealed structures and the expansion of the exposed siphon wall (Fig. 1e).

To distinguish between these mechanisms, we coated the contracted posterior siphon with fluorescent dye and manually extended it using forceps (Fig. 1f). If extension involves unfolding of a concealed region, the newly exposed surface would remain unlabeled. In contrast, if extension occurs exclusively through expansion of the exposed siphon wall, the fluorescent label would be expected to remain distributed across the extended surface.

Following manual extension, a large non-fluorescent region became exposed in the distal portion of the posterior siphon (Fig. 1f). The length of this region corresponded to approximately half of the larval body length (median non-fluorescent region-to-body length ratio = 0.53, IQR = 0.481–0.589, *n* = 6; Fig. 1g). These observations provide experimental support for the previously proposed folding– unfolding mechanism, in which concealed portions of the posterior siphon become exposed during extension. Although expansion of the siphon wall may also contribute, unfolding appears to account for a substantial component of posterior siphon elongation.

### The posterior siphon contains an extensive array of transverse muscle fibers

Given the remarkable extensibility of the posterior siphon, we investigated its muscular organization using phalloidin staining to identify structural features potentially associated with siphon extension. Phalloidin staining revealed numerous transverse muscle fibers distributed throughout the posterior siphon of *H. virgatus*. Quantification of fiber orientation showed that most muscle fibers were oriented approximately perpendicular to the longitudinal axis of the siphon, demonstrating a strong bias toward transverse orientations (Fig. 2a, a″). In contrast, relatively few transverse muscle fibers were observed in abdominal segments A8 of *Episyrphus* sp. and *D. melanogaster* (Fig. 2b, c). Although technical difficulties in completely dissecting the posterior siphon prevented us from determining the total number of muscle fibers, more than 200 transverse muscle fibers intersecting a single longitudinal muscle were observed on one side of the posterior siphon in each *H. virgatus* specimen examined (*n* = 4). By comparison, no more than 20 such fibers were observed in abdominal segment A8 of *Episyrphus* sp. (*n* = 3) and *D. melanogaster* (*n* = 3).

**Figure 2.**
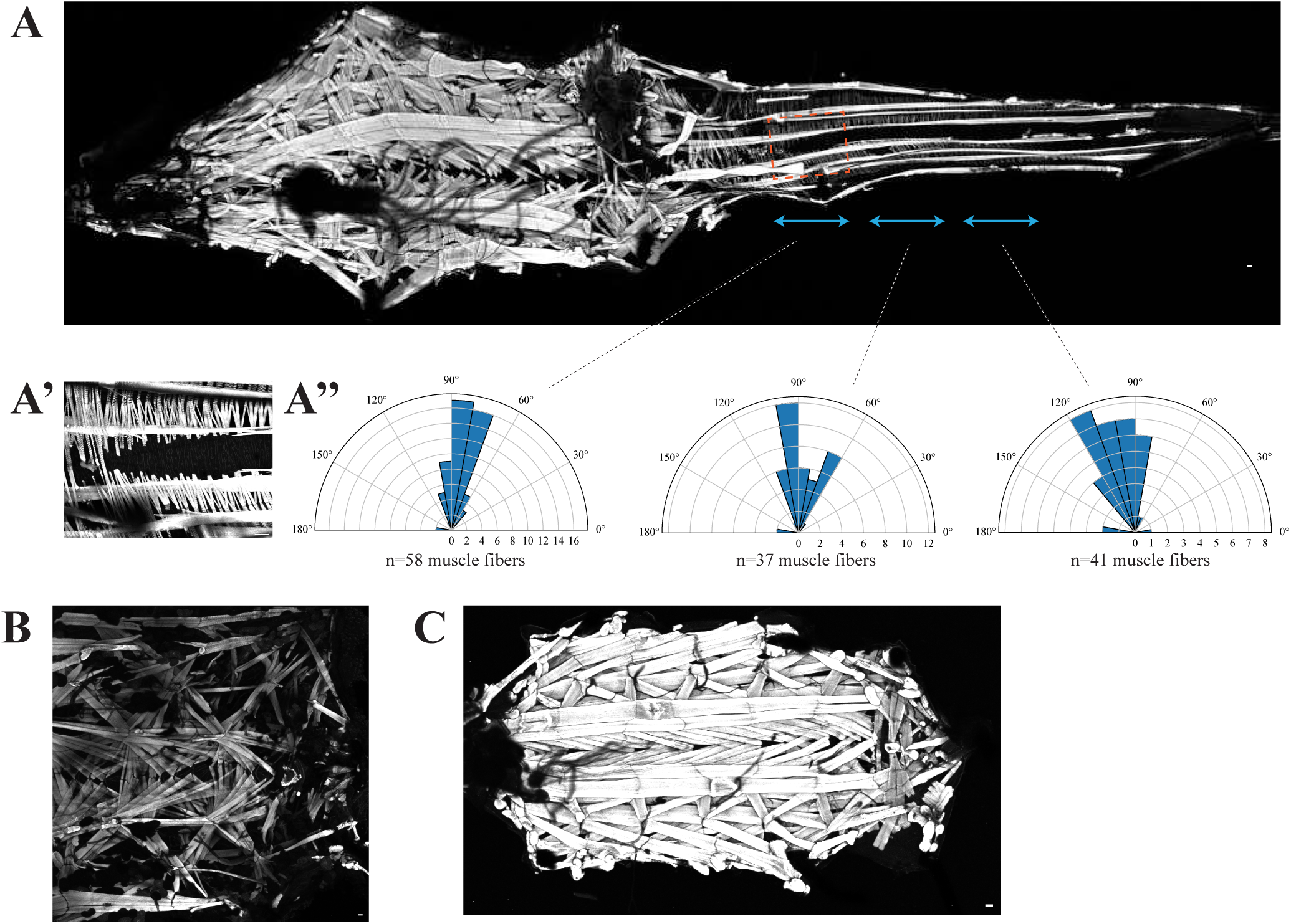
Structural basis of posterior siphon extensibility in *Helophilus virgatus*. **(a)** Representative phalloidin staining of an *H. virgatus* larval fillet preparation (*n* = 4). Anterior is to the left. **(a′)** Enlarged view of the boxed region in the posterior siphon. **(a″)** Rose plots showing muscle fiber orientations in the proximal (left), middle (center), and distal (right) regions of the posterior siphon, indicated by sky-blue arrows in **(a)**. Fiber orientation was measured relative to the siphon axis (0° = parallel to the siphon axis). **(b)** Representative phalloidin staining of the posterior body region (approximately abdominal segments A5–A8) of an *Episyrphus* sp. larva (*n* = 3). **(c)** Representative whole-body phalloidin staining of a *D. melanogaster* larva (*n* = 3). Scale bars, 50 μm.

These observations indicate that the extensible posterior siphon of *H. virgatus* contains a striking enrichment of transversely oriented muscle fibers, consistent with the hypothesis that these muscles contribute to siphon extension.

### *Helophilus* larvae exhibit siphon-mediated near-surface locomotion

Some aquatic insects swim beneath the water surface, whereas others float and move along the air–water interface [22,23]. We investigated the near-surface locomotion of *H. virgatus* larvae and its associated kinematics. We first observed that *H. virgatus* larvae remained afloat even when their posterior spiracles were submerged beneath the water surface, whereas *D. melanogaster* larvae rapidly sank under the same conditions. This observation suggested a difference in the ability of the two species to remain afloat. Because air-filled tracheal structures may contribute to buoyancy in aquatic insect larvae [17], we compared the relative width of the dorsal tracheal trunk, a major longitudinal component of the tracheal system, between the two species (Fig. 3a). The dorsal tracheal trunk-to-body width ratio was significantly larger in *H. virgatus* than in *D. melanogaster* (median = 0.21, IQR = 0.17–0.22, *n* = 8, versus median = 0.046, IQR = 0.039–0.049, *n* = 8; Fig. 3b). This difference suggests that the enlarged tracheal system may contribute to the ability of *H. virgatus* larvae to remain afloat, although its contribution to buoyancy was not directly measured.

**Figure 3.**
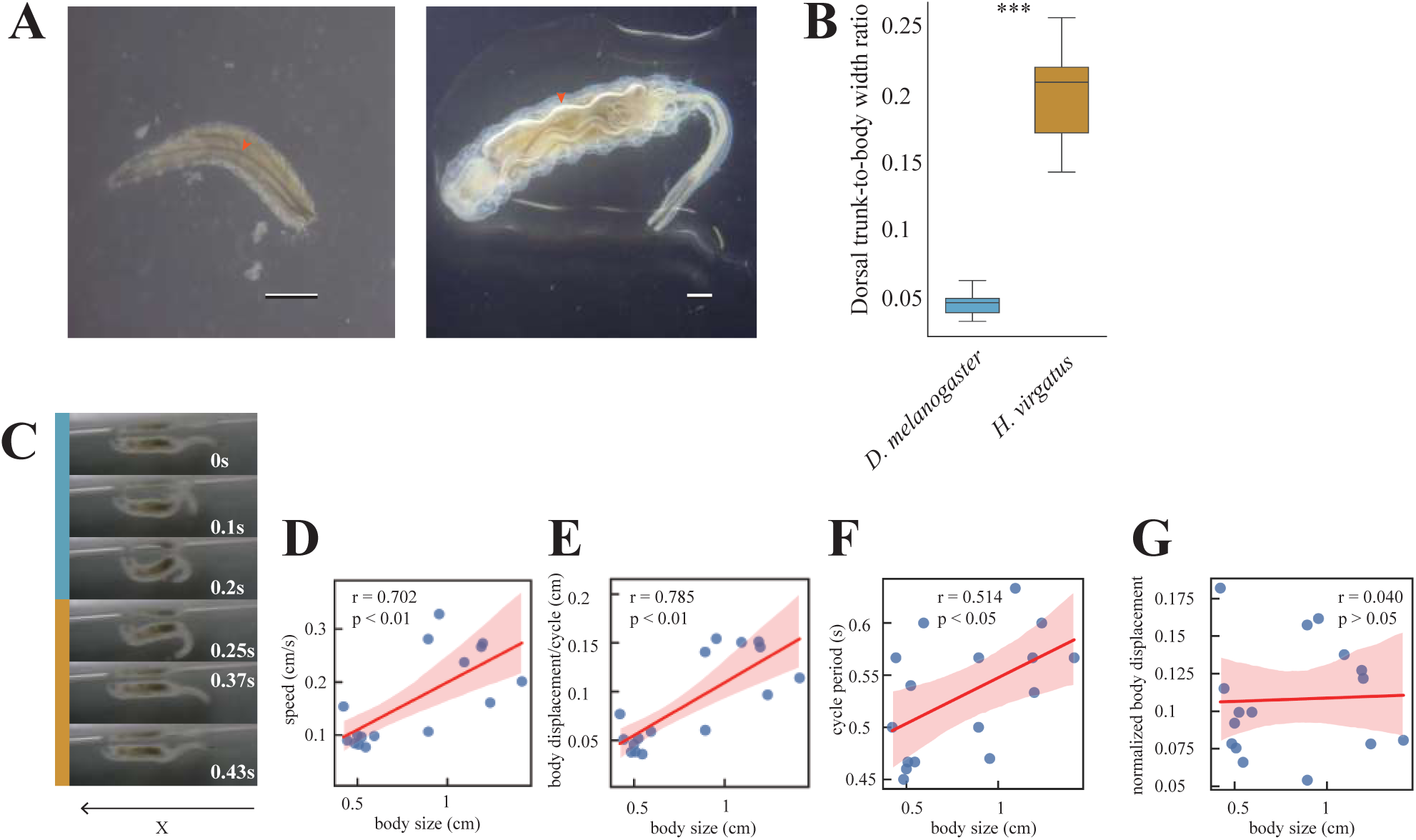
Morphological characteristics and kinematics of near-surface locomotion in *Helophilus* larvae. **(a)** Representative dorsal views of *D. melanogaster* (left) and *H. virgatus* (right) larvae on agar plates. Dorsal trunks are indicated by orange arrowheads. Scale bars 1 mm. **(b)** Box plot showing the dorsal trunk-to-body width ratio in *D. melanogaster* and *H. virgatus*. N= 8 each. \*\**P* < 0.001, Mann–Whitney U test. **(c)** Time-lapse images showing near-surface locomotion of an *H. virgatus* larva driven by rowing movements of the posterior siphon. The forward stroke (0–0.25 s, sky blue) is followed by the backward stroke (0.25–0.466 s, orange). **(d–g)** Relationships between body size and swimming speed (d; cm s⁻¹, Pearson’s *r* = 0.702, *P* < 0.01), body displacement per rowing cycle (e; cm, *r* = 0.785, *P* < 0.01), rowing cycle period (f; s, *r* = 0.514, *P* < 0.05), and normalized body displacement per rowing cycle (g; *r* = 0.040). For panels (d–g), *n* = 16 larvae. Red lines indicate linear regression fits, and shaded areas represent 95% confidence intervals.

Under resting conditions, larvae typically maintain the posterior spiracles at the air–water interface for respiration. Before initiating locomotion, however, larvae first submerged the posterior siphon beneath the water surface while remaining afloat (Movie S2). During locomotion, larvae were typically oriented with the ventral side facing upward and repeatedly depressed the submerged posterior siphon before elevating it, in an asymmetric rowing motion associated with forward movement (Fig. 3c; Movie S2-3).

To determine whether near-surface locomotion scales with body size, we examined the relationships between body size and locomotor performance. Body size was positively correlated with swimming speed (*r* = 0.702, *P* < 0.01), body displacement per rowing cycle (*r* = 0.785, *P* < 0.01), and rowing cycle period (*r* = 0.514, *P* < 0.05), but not with normalized displacement per cycle (*r* = 0.040, *P* > 0.05; n = 16 larvae; Fig. 3d–g). Thus, larger larvae tended to travel farther during each rowing cycle but also exhibited longer cycle periods, whereas displacement normalized to body size showed no detectable relationship with body size.

### Asymmetric siphon strokes are associated with forward body displacement

These observations prompted us to examine how the movements and deformation of the posterior siphon relate to forward body displacement during individual rowing cycles. We tracked the trajectory of the posterior siphon tip relative to its base and the displacement of the midbody point, and quantified the base–tip angle (θ) and siphon straightness throughout each rowing cycle (Fig. 4a, a′).

**Figure 4.**
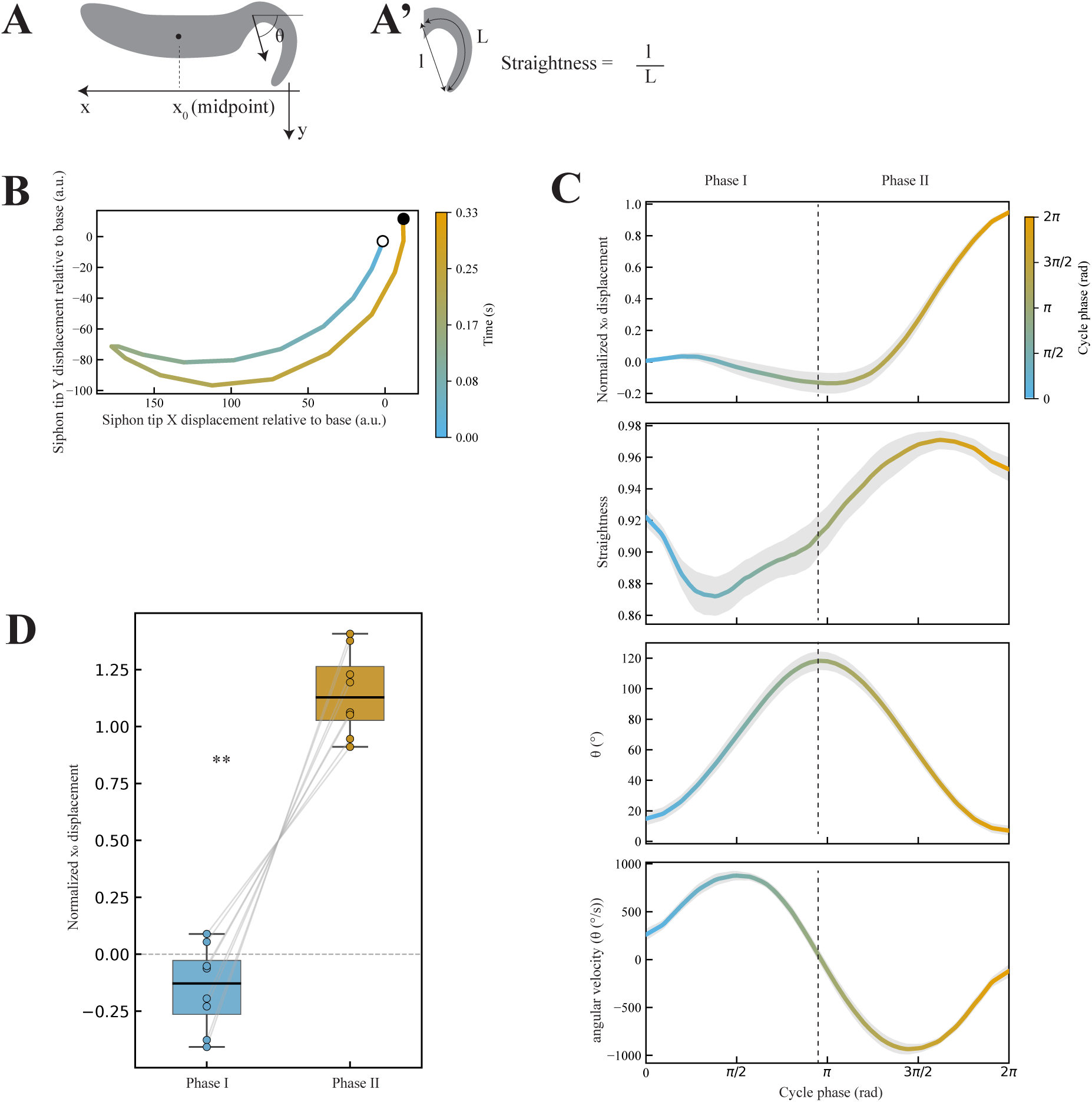
Asymmetric body displacement during forward and backward strokes of the posterior siphon in *Helophilus* larvae. **(a)** Schematic illustrating the definitions of (x_0_), the horizontal coordinate of the midpoint between the siphon base and the anterior tip of the larval body, and θ, the angle between the horizontal water surface and the line connecting the siphon base to its tip. The positive x- and y-directions are defined as leftward and downward, respectively. **(a′)** Schematic illustrating siphon straightness, defined as the ratio of the straight-line distance ((l)) to the contour length ((L)) between the siphon base and tip ((l/L)). **(b)** Representative trajectory of the posterior siphon tip relative to its base during a single rowing cycle. The initial tip position was set to the origin (0, 0). Colour indicates elapsed time, progressing from cyan to orange. Open and filled circles indicate the beginning and end of the rowing cycle, respectively. **(c)** Changes in normalized (x_0_) displacement, siphon straightness, base–tip angle (θ), and angular velocity over the rowing cycle (from top to bottom; *n* = 8 larvae). Time was normalized to a cycle phase ranging from 0 to 2π, and (x_0_) displacement was normalized to the net displacement over each cycle, such that the values at the beginning and end were 0 and 1, respectively. Solid lines indicate the mean, and shaded regions represent the standard error of the mean (SEM). The dashed vertical line approximately indicates the transition between Phase I and Phase II. **(d)** Box plots comparing normalized (x_0_) displacement during Phase I and Phase II (*n* = 8 larvae). The two phases were defined for each larva by the transition of angular velocity from positive to negative. Grey lines connect measurements from the same larva. **P < 0.01, two-sided Wilcoxon signed-rank test.

Each rowing cycle was divided into two phases based on the reversal of siphon angular velocity from positive to negative: a forward stroke (Phase I), during which the siphon tip moved forward, and a backward stroke (Phase II), during which it moved backward (Fig. 4b, c; Figure supplement 1a–h; *n* = 8 larvae). Siphon straightness decreased during the early part of Phase I, indicating increased curvature, and subsequently increased as the siphon returned towards a straighter configuration (Fig. 4c; Figure supplement 1a–h).

Analysis of midbody displacement revealed a pronounced asymmetry between the two phases. The midbody point showed little net forward displacement or slight backward movement during Phase I, whereas most forward displacement occurred during Phase II (Fig. 4c, d; two-sided Wilcoxon signed-rank test, *P* < 0.01; *n* = 8 larvae). During the backward stroke, both the anterior end of the body and the siphon base also moved forward, consistent with forward translation of the larva rather than merely a change in body shape (Movie S3).

Finally, we quantified kinematic rowing effectiveness as the absolute net displacement of the midbody point along the x-axis divided by the total path length travelled by the posterior siphon tip relative to its base during a single rowing cycle. The median kinematic rowing effectiveness was 0.073 (interquartile range = 0.043–0.074; *n* = 5 larvae; Figure supplement 1i). These findings demonstrate that posterior siphon rowing is characterized by asymmetric deformation and body displacement across the two phases of the movement cycle.

## DISCUSSION

In this study, we investigated the structural and functional specializations of the extensible posterior respiratory siphon in larvae of *Helophilus virgatus*. Fluorescence labeling provided experimental evidence supporting the previously proposed unfolding of concealed anal structures during siphon extension [16]. Phalloidin staining further revealed an extensive array of transversely oriented muscle fibers within the posterior siphon, a feature that was less prominent in the corresponding posterior regions of *Episyrphus* sp. and *Drosophila melanogaster*. Behavioral and kinematic analyses revealed a previously undescribed mode of near-surface locomotion involving asymmetric rowing movements of the posterior respiratory siphon, with most forward body displacement occurring during the backward stroke. Together, these findings demonstrate that a highly extensible respiratory structure can also participate in locomotion, highlighting the multifunctionality of specialized body structures in aquatic environments.

The folding architecture of the posterior siphon may provide several mechanical advantages. Folding allows substantial extension while maintaining a compact resting configuration, potentially reducing the spatial constraints associated with a permanently elongated structure. Similar folding-based deployment strategies occur in diverse organisms, including the hyperextensible neck of the ciliate *Lacrymaria olor*, which extends through a curved-crease origami mechanism [24], the foldable hindwings of rove beetles [25], and the deployable helmets of treehoppers [26]. These examples illustrate how folding-based architectures can accommodate large changes in shape while maintaining compact configurations. Although the precise geometry of the siphon folds remains to be determined, our findings provide experimental support for unfolding as an important component of posterior siphon extension.

Structures that evolved in association with one function can subsequently acquire additional functions during evolution [27]. Our observations show that the posterior respiratory siphon of *H. virgatus* is involved in both respiration and near-surface locomotion. One possible evolutionary scenario is that elongation of the posterior respiratory structure initially facilitated access to atmospheric oxygen while the larval body remained submerged, with its subsequent use in rowing representing an additional locomotor function. This would be consistent with evolutionary co-option, although the sequence in which these functions evolved remains unknown. Comparative analyses across eristaline species differing in siphon morphology and locomotor behavior could help determine whether the evolution of an elongated respiratory siphon preceded its recruitment for rowing-based locomotion.

Rowing-based locomotion involving distinct power and recovery strokes has been documented in animals moving at the air–water interface, including water striders [28,29] and fishing spiders [30]. Unlike these animals, which use their legs for propulsion, *H. virgatus* larvae employ a respiratory siphon during near-surface locomotion. Our kinematic analyses revealed pronounced asymmetry between the two phases of the rowing cycle: most forward body displacement occurred during the backward stroke, whereas the forward stroke produced little net forward displacement. Siphon straightness also varied over the cycle, decreasing during the early forward stroke and subsequently increasing. One possible explanation is that changes in siphon curvature alter its hydrodynamic resistance, allowing the backward stroke to generate greater forward thrust. Under this interpretation, the backward stroke may function as a power stroke, whereas the forward stroke may serve primarily as a recovery stroke. However, direct measurements of fluid motion and hydrodynamic forces are needed to establish the mechanism of propulsion and determine the contribution of siphon deformation.

Together, our findings reveal how an extensible respiratory structure can participate in a distinct locomotor behavior. The combination of a folding architecture, specialized musculature and asymmetric rowing movements illustrates how morphological specializations associated with respiration can provide the basis for additional mechanical functions in aquatic environments.

## ACKNOWLEDGEMENTS

We thank the Kyoto Drosophila Stock Center for providing a *Drosophila* strain. We are grateful to the members of the Nose laboratory for fly food preparation, as well as helpful discussions and feedback. This work was supported by the Japan Society for the Promotion of Science (grant no. 25K09722 to T.M.) and the Toyota Foundation (grant no. D24-HS-0144 to T.M.).

## FIGURE SUPPLEMENTS

**Figure supplement.**
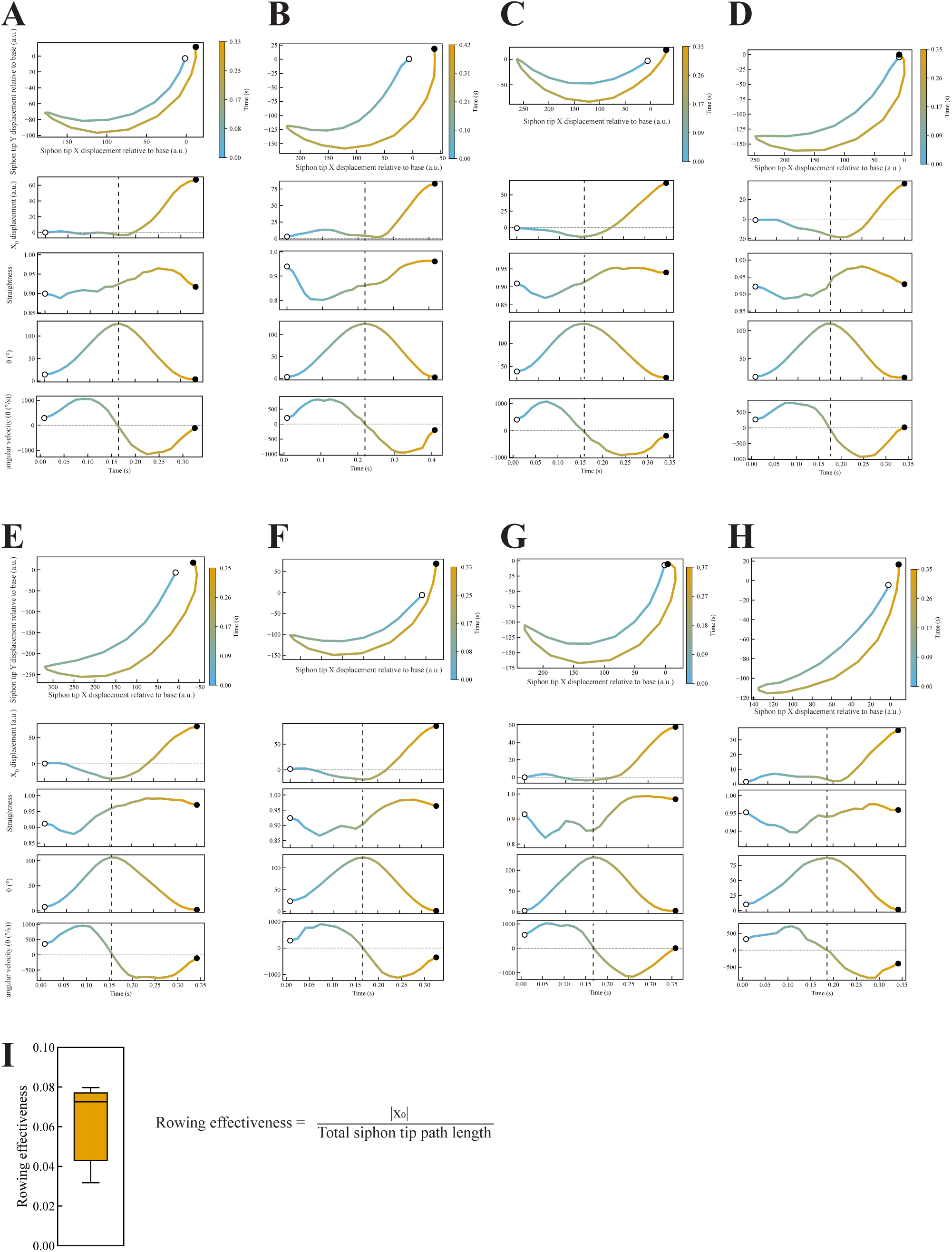
Individual trajectories of posterior siphon rowing movements and kinematic rowing effectiveness in *Helophilus* larvae. **(a–h)** Individual kinematics of posterior siphon rowing during a single cycle. From top to bottom, each panel shows the trajectory of the posterior siphon tip relative to its base, (x_0_) displacement, siphon straightness, base–tip angle (θ), and angular velocity as a function of time (s). Siphon tip coordinates were calculated relative to the siphon base, with the initial tip position set to the origin (0, 0). Colour indicates elapsed time, progressing from cyan to orange. Open and filled circles indicate the beginning and end of the rowing cycle, respectively. Vertical dashed lines indicate the transition between Phase I and Phase II, defined as the point at which angular velocity changes from positive to negative. **(i)** Box plot showing kinematic rowing effectiveness, defined as the absolute net displacement of (x_0_) along the x-axis divided by the total path length travelled by the posterior siphon tip relative to its base during a single rowing cycle (*n* = 5 larvae).

## SUPPLEMENTARY FILES

**Movie S1. Posterior siphon elongation for respiration in a *H. virgatus* larva.**

An *H. virgatus* larva extends its posterior siphon to maintain contact with the air-water interface for respiration. Playback speed: 1×.

**Movie S2. Submergence of the posterior spiracles preceding near-surface locomotion in a *H. virgatus* larva.**

An *H. virgatus* larva submerges its posterior spiracles beneath the air–water interface before propelling itself just beneath the interface by rowing with its posterior siphon. Playback speed: 1×.

**Movie S3. Near-surface locomotion of a *H. virgatus* larva.**

An *H. virgatus* larva propels itself just beneath the air–water interface by rowing with its posterior siphon. Playback speed: 0.5×.

